# Automating RNA-Ligand Interaction Modeling via a Self-Improving LLM Agent

**DOI:** 10.1101/2025.09.11.675747

**Authors:** Zhejun Kuang, Yunkai Li, Yihang Bao, Shengyang Zhou, Zeqi Dong, Weidi Wang, Guan Ning Lin, Han Wang, Zhe Liu

## Abstract

Precise modeling of RNA-ligand interactions is essential for understanding RNA functionality and designing RNA-targeted therapeutics. Current computational approaches largely focus on predicting discrete binding sites, limiting their applicability to complex RNA regions that may harbor multiple or diffuse ligand binding motifs. Here, we present RLAgent, an interactive agent framework designed to predict ligand interactions at the RNA region level, enabling higher-resolution and more flexible modeling than conventional site-centric approaches. RLAgent reframes the RNA-ligand prediction workflow as a dialogue-driven process. Through a natural language interface, users can interactively configure modeling preferences without writing code. A locally hosted large language model (LLM) acts as the core orchestration agent, automating all key components of the modeling pipeline, including data validation, feature encoding, model training, evaluation, and visualization. This agent-based design lowers technical barriers and enhances reproducibility, making RNA-ligand prediction more accessible for both computational and experimental researchers.

## I. Introduction

Understanding and predicting RNA-ligand interactions is a key step toward deciphering RNA function and developing RNA-targeted therapeutics[1-4]. These interactions play essential roles in processes such as translation regulation, riboswitch activation, and small-molecule drug targeting[5-7]. Despite advances in RNA structural biology and ligand screening technologies, computational models for RNA-ligand binding remain limited in scope, particularly in terms of resolution and accessibility[8-10].

Most existing prediction models focus on either full-length RNA structures or localized binding sites[1, 8, 11-15]. While these approaches have demonstrated utility in identifying key interaction points, they often fail to capture the complexity of biologically relevant RNA segments based solely on RNA sequences, which may contain multiple, distributed, or structurally ambiguous binding motifs. This restricts their flexibility in modeling interactions over short or non-canonical regions of interest, particularly in therapeutic or synthetic contexts where RNA editing or motif engineering is applied.

Although RNA-ligand datasets are becoming increasingly available, constructing effective prediction models remains challenging in practice. Existing workflows typically require users to complete multiple steps manually[11, 13, 15], including data preprocessing, model selection, and performance evaluation. These steps are often implemented through scripting interfaces and command-line tools. This not only introduces technical barriers, especially for biologists or non-specialists, but also limits the reproducibility and scalability of RNA-targeted modeling pipelines.

Recent developments in large language models (LLMs) have enabled systems that can interpret natural language instructions and generate executable code across a wide range of tasks[16-20]. This capability presents an opportunity to simplify the modeling process by reducing the need for manual scripting and enabling more accessible, flexible pipeline construction. However, these capabilities have not yet been widely applied to automate RNA-ligand interaction modeling in a user-friendly and domain-aware manner.

To address these limitations, we introduce RLAgent, an interactive agent framework designed to support RNA region-ligand interaction modeling based on RNA sequences and ligand property. Unlike existing approaches that focus on isolated binding sites or transcript-level properties, RLAgent operates at the region level, enabling more flexible and informative modeling of RNA segments that may contain multiple or diffuse ligand binding motifs.

RLAgent integrates large language model capabilities with optional retrieval-augmented generation (RAG), optional RNA foundation model-based feature encoding, automated debugging, and multimodal feature integration, forming a coherent, end-to-end pipeline. By allowing researchers to specify tasks, adjust workflows, and refine models entirely through dialogue, RLAgent removes the technical barriers imposed by traditional scripting-based approaches, thereby enhancing accessibility for non-specialists while preserving methodological rigor. The source code of the pre-processing, modeling and validation processes are freely available on GitHub (https://github.com/Liuzhe30/RLAgent).

## II. Methods

### A. RLAgent Workflow

RLAgent is an interactive agent framework designed to enable fully natural language-driven execution of data processing, model construction, training, validation, and visualization for RNA-ligand region-level interaction prediction (Figure 1). The framework integrates large language model (LLM) reasoning with optional retrieval-augmented generation (RAG), automated self-debugging, and RNA foundation model-based feature encoding, thereby forming an end-to-end system capable of translating user intentions into executable analytical workflows. This design substantially reduces the technical barrier of computational modeling, particularly for researchers without programming expertise, while ensuring methodological reproducibility through standardized, machine-generated code.

**Fig. 1.**
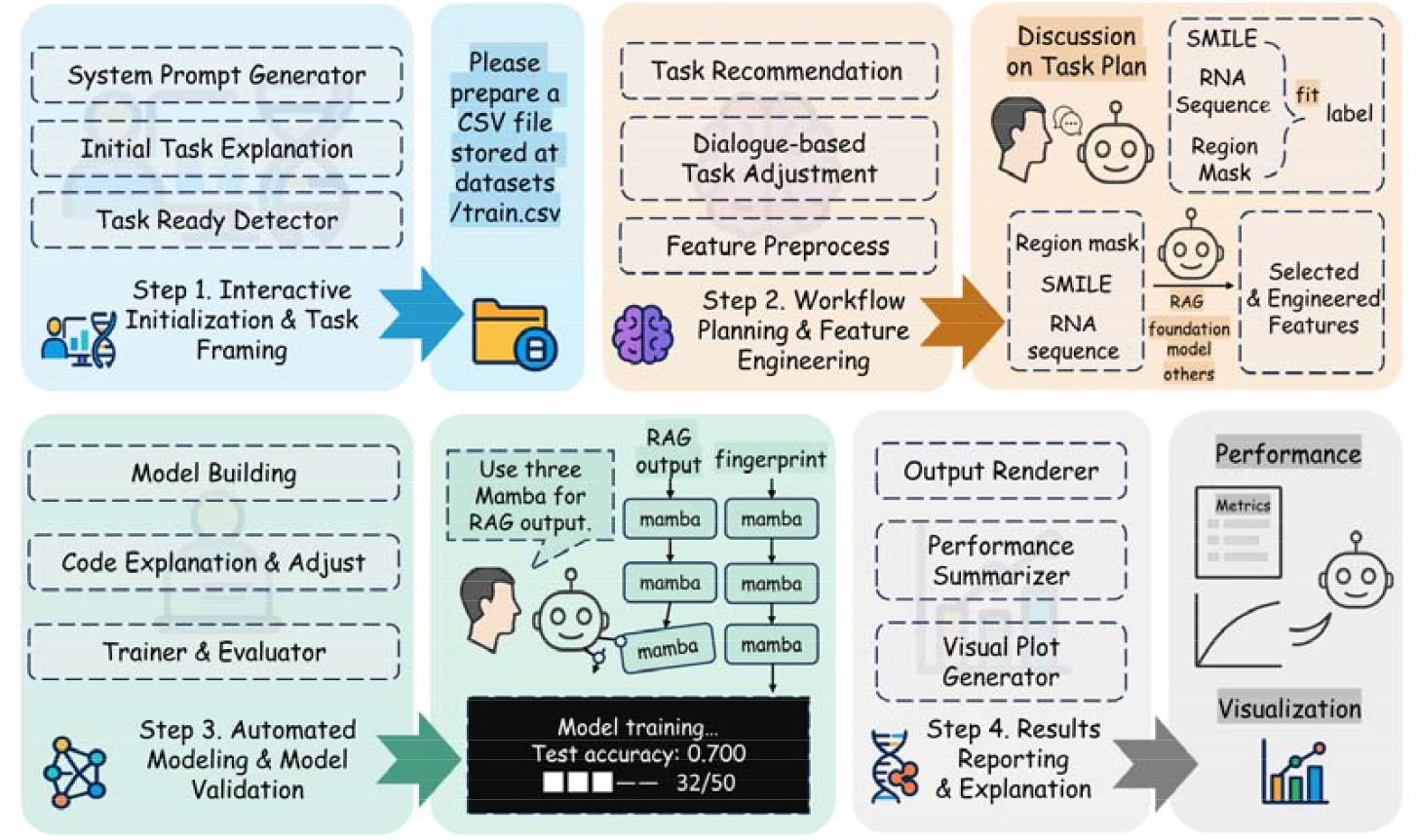
Overview of RLAgent. RLAgent is designed as an interactive agent system that transforms natural language instructions into fully executable RNA region-ligand prediction workflows.

#### 1) Interactive initialization and task framing

At the start of each session, RLAgent conducts a dialogue-based initialization process that interprets user objectives and constructs a structured workflow plan. The system first issues system-level prompts and provides an initial summary of the modeling task, including its biological context, required data modalities, and expected outputs. It then examines whether the input data meet readiness criteria, such as valid sequence formatting, ligand identifiers, and region mask definitions, and guides the user in preparing a standardized dataset. Each entry in this dataset contains an RNA sequence, a corresponding ligand represented by its SMILES string, and a binary mask specifying the region of interest. This preparatory step, executed through conversational prompts, ensures data consistency and completeness prior to model construction. The entire interaction is supported by DeepSeek-V1-70B, which enables contextual understanding of user instructions and precise generation of task-specific guidance.

#### 2) Workflow planning and feature engineering

The agent recommends task configurations, supports dialogue-based adjustments, and preprocesses features. SMILES representations of ligands, RNA sequences, and region masks are combined with optional RAG outputs and optional embeddings from RNA foundation model (RNA-FM)[21] to construct a set of selected engineered features tailored to the modeling objective.

For users who activate the optional RAG module, RLAgent retrieves external biochemical knowledge from an ontology-based knowledge graph. These retrieved entities are transformed into auxiliary numerical features, thereby enriching the model input space with contextual molecular information. The resulting multimodal feature vectors are standardized and concatenated into a unified format for downstream modeling.

#### 3) Automated modeling and model validation

Following feature assembly, RLAgent autonomously generates the model training pipeline, including data partitioning, architecture construction, optimization setup, and evaluation metrics. The generated code is continuously reviewed by a built-in self-iterative debugging module, which recursively detects and corrects execution errors or inconsistencies between generated scripts and user-defined requirements.

RLAgent supports both classical machine learning (e.g., Random Forest, XGBoost, LightGBM) and deep learning architectures (e.g., LSTM, Transformer, and Mamba). Deep architectures are particularly advantageous for modeling complex dependencies between RNA sequence patterns and ligand properties. The system monitors training progress in real time, reporting quantitative indicators such as accuracy, precision, recall, F1-score, and the area under the ROC curve (AUC). In addition to standard evaluation, the agent records the number of debugging iterations required for successful execution, providing an implicit measure of workflow robustness and model reproducibility.

#### 4) Results reporting and explanation

Upon completion of model training and evaluation, RLAgent automatically compiles a comprehensive report that integrates both quantitative and visual outputs. This includes numerical performance metrics, confusion matrices, receiver operating characteristic (ROC) curves, and other diagnostic plots. The report also provides LLM-generated textual explanations of model behavior and performance interpretation in plain language, helping users understand the analytical outcomes.

All results are generated in standardized formats suitable for downstream comparison, publication, or integration into laboratory data systems. This reporting stage completes the end-to-end pipeline, transforming a natural language query into a fully reproducible RNA-ligand interaction model accompanied by transparent, interpretable documentation.

### B) Self-iterative Debugging Process

All code generation tasks for RLAgent utilize self-iterative debugging to ensure code executability. For each requirement in the feature engineering, modeling, and results reporting stages, the RLAgent uses LLM to initialize a piece of code and employs automatic code correction methods to refine the code’s content and executability. The correction methods will adjust the code based on requirements and potential bugs and revalidating the corrected code to ultimately produce code that have content aligns with the requirements and is free of errors.

#### Algorithm 1

workflow of automatic code correction

**Input:** Code snippet *code*; user requirement *query*

**Output:** Corrected executable code *code*

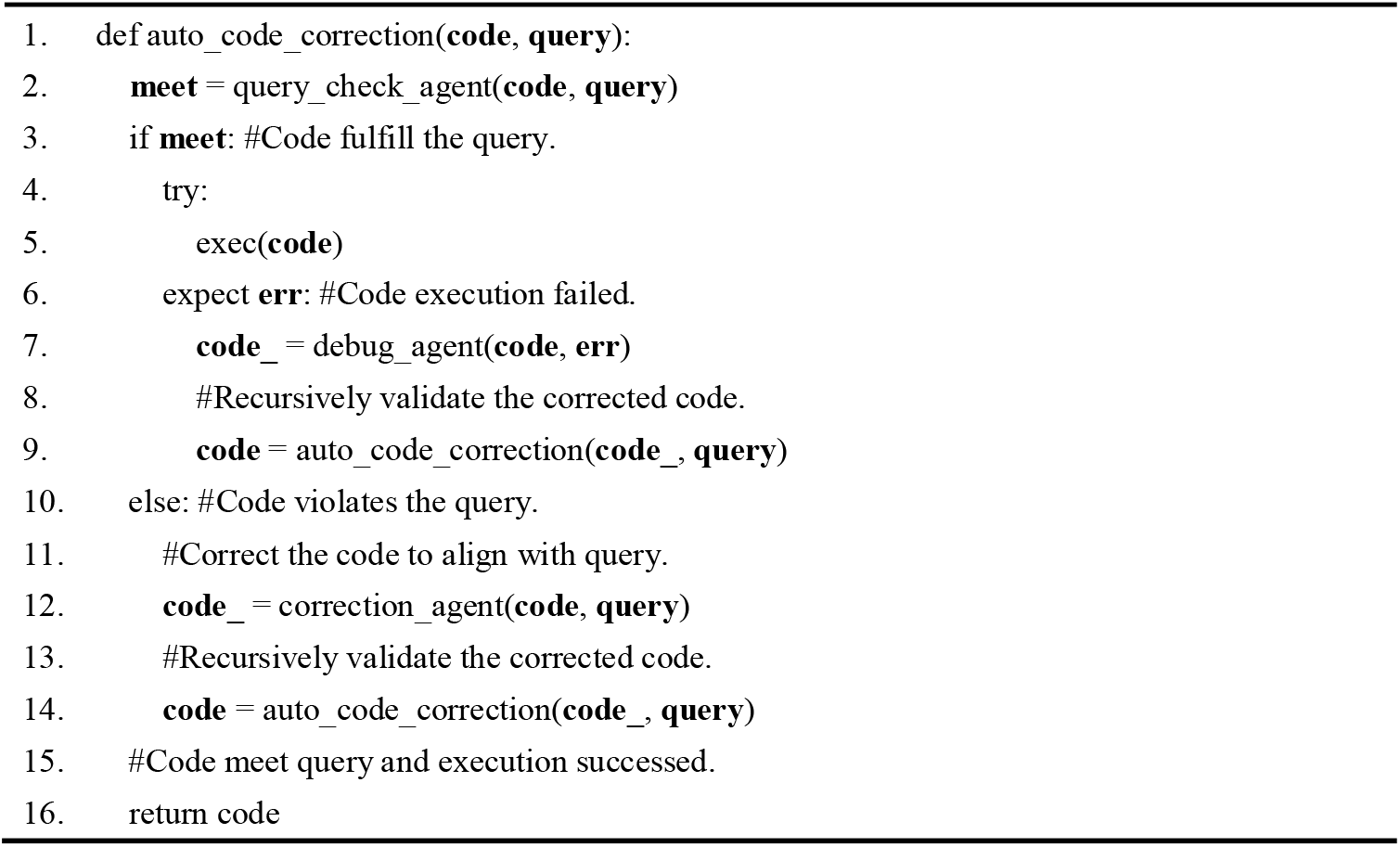

### C. Datasets and Task Definition

We formulate RNA region-ligand interaction prediction as a binary classification task. Given an RNA sequence with a specified region and a ligand compound, the model determines whether the ligand binds to this region within the RNA molecule, aiming to capture the fine-grained pairing preferences between chemical ligands and specific RNA segments.

Positive samples were collected from RNALigands [22], a curated database of experimentally verified RNA-ligand complexes (Figure 2). The datasets used in this study were derived from the Protein Data Bank (PDB). Each entry provides precise region-level binding information, and region-level annotations were retrieved and aligned to corresponding RNA sequences using the RCSB PDB FASTA API.

**Fig. 2.**
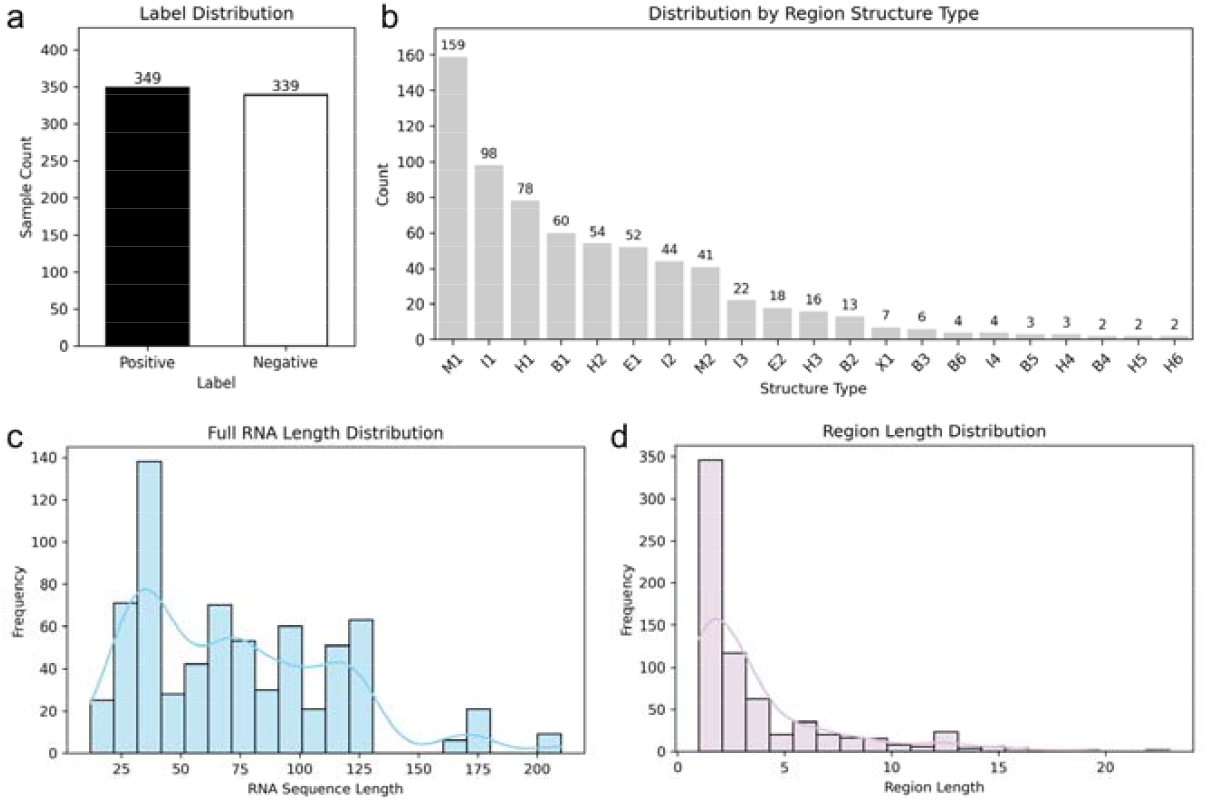
Dataset analysis of RNA region-ligand prediction task in this study.

For cases where a single region annotation corresponds to multiple disjoint binding subsequences, each fragment was treated as an independent training sample. In cases where the region comprised a single nucleotide, we adopted a left-most match heuristic to localize the fragment within the full sequence [23]. Region annotations that could not be reliably localized, or those missing ligand or region information, were excluded. For each positive RNA region, we randomly paired it with a ligand not observed to bind that region in any known structure, assuming no interaction occurs. The selected ligand differs from the original positive pair, and the overall dataset maintains a 1:1 ratio of positive to negative samples.

Ligands were annotated with structural fingerprints and SMILES representations via the PubChem database[24]. Ligands without valid annotations were excluded from the dataset. After preprocessing and filtering, the final dataset contains 349 positive and 339 negative samples, as shown in Fig. 2.

### D. LLM Backends for RLAgent

RLAgent was powered by multiple LLMs with varying parameter scales and deployment modes. GPT-4o (200B, released May 2024) was accessed through the MetaChat API (https://www.metachat.work/#/chat), while all other models were locally deployed, including DeepSeek-R1:70B (January 2025), deepseek-v2:16B (May 2024), deepseek-r1:7B (January 2025), and LLaMA3.2:3B (January 2024).

### E. Feature Encoding and Model Options

To support region-level RNA-ligand interaction modeling, we employ a hybrid representation strategy that combines RNA sequence embeddings with ligand molecular fingerprints. For RNA input, we use one-hot encoding, with optional incorporation of embeddings derived from the RNA foundation model (RNA-FM)[21] and optional integration of retrieval-augmented generation (RAG). The RAG component retrieves external domain knowledge from the ontology-based knowledge graph for representing interactions involving RNA molecules[25], in which retained information includes entities that can be located in Google Scholar using the search terms “Protein”, “GO”, “Disease”, “miRNA”, “Pathway”, and “Phenotype”. Each region is located within the full-length sequence via a binary mask. For ligand input, we use fingerprints computed from SMILES annotations retrieved via PubChem.

RLAgent supports multiple model types to accommodate diverse data scales and user preferences. For classical machine learning, users may choose from Random Forest, Naïve Bayes, XGBoost, or LightGBM. These models are fast to train and interpretable, making them suitable for small or imbalanced datasets. For more expressive modeling, the framework includes neural architectures such as long short-term memory (LSTM) networks, transformer-style self-attention encoders, and the Mamba sequence model[26], which has recently shown strong performance on token-level classification tasks. All models are trained on the concatenated RNA-ligand representations, and model selection is interactively guided through natural language dialogue.

### D. Training and Evaluation Details

Under fixed parameters and 50 training iterations without using early stopping, the dataset was randomly split into 80% for training and 20% for testing. Classical machine learning models were implemented using scikit-learn, while deep learning models were developed with PyTorch. The deep learning models were trained for 50 epochs. Each deep learning model encoded RNA and ligand features separately through two feedforward layers, concatenated the resulting vectors, and passed them through two additional linear layers to produce the final prediction.

For performance evaluation, we reported standard binary classification metrics in this study, including accuracy, precision, recall, F1-score, and the area under the ROC curve (AUC). In addition, because RLAgent employs an automated debugging mechanism, workflows could often be executed to completion even when generated code did not strictly match user instructions. To capture this aspect, we also recorded the number of cases requiring manual inspection to verify code conformity across repeated runs.

## III Results

### A. Robustness Analysis of Underlying LLMs

We assessed the robustness of different LLM backends in RLAgent by running eight complete workflows for each model. At every interaction step, success was recorded only if the LLM produced code that fully conformed to user requirements within 200 automated debugging iterations. Failures included non-convergence within the debugging budget, generated is not exactly as expected and premature termination due to formatting errors. This metric therefore reflects the practical usability of code outputs rather than execution success alone.

As summarized in Table 1, GPT-4o exhibited the highest robustness, achieving 100% success in feature extraction, pooling, and code assembly, and 87% in multilayer perceptron implementation. DeepSeek-R1:70B also performed reliably, with moderately lower but consistent success across modules. In contrast, models with fewer than 20B parameters failed to complete any step successfully under the defined criteria, indicating insufficient reliability for integration into RLAgent without substantial error correction. These results suggest that larger-scale LLMs are substantially more robust for domain-specific code generation, reducing the dependency on debugging and enhancing the reliability of autonomous workflows. This observation also provides a practical guideline for selecting LLM backends in future agent-based systems, where model scale and robustness must be balanced against efficiency and deployment constraints.

**Table 1.** Values represent the proportion of successful cases across eight complete RLAgent workflows. A step is counted as successful only if the generated code fully satisfied the user’s requirements within a maximum of 200 automated debugging iterations.

| Model | Parameters | Release Date | Success Rate - Feature | Success Rate - | Success Rate - MLP | Success Rate - Code |
| --- | --- | --- | --- | --- | --- | --- |

|  |  |  | Extraction | Pooling |  | Assembly |
| --- | --- | --- | --- | --- | --- | --- |
| GPT-4o | 200B | 2024.05 | 100% | 100% | 87% | 100% |
| DeepSeek-R1:70B | 70B | 2025.01 | 87% | 87% | 75% | 87% |
| DeepSeek-V2:16B | 16B | 2024.05 | 0% | 0% | 0% | 0% |
| DeepSeek-R1:7B | 7B | 2025.01 | 0% | 0% | 0% | 0% |
| LLaMA3.2:latest | 3B | 2024.01 | 0% | 0% | 0% | 0% |

### B. Evaluating the Functional Contributions of Querying, Debugging, and Generation Strategies in RLAgent

To investigate the contribution of individual components within RLAgent, we performed an ablation study on locally deployed DeepSeek-R1:70B, repeating each configuration across eight full workflow runs (Table 2). The baseline setting reflects the standard interactive workflow, while three alternative configurations were tested: removal of query validation, removal of debugging, and single-pass code generation.

**Table 2.**
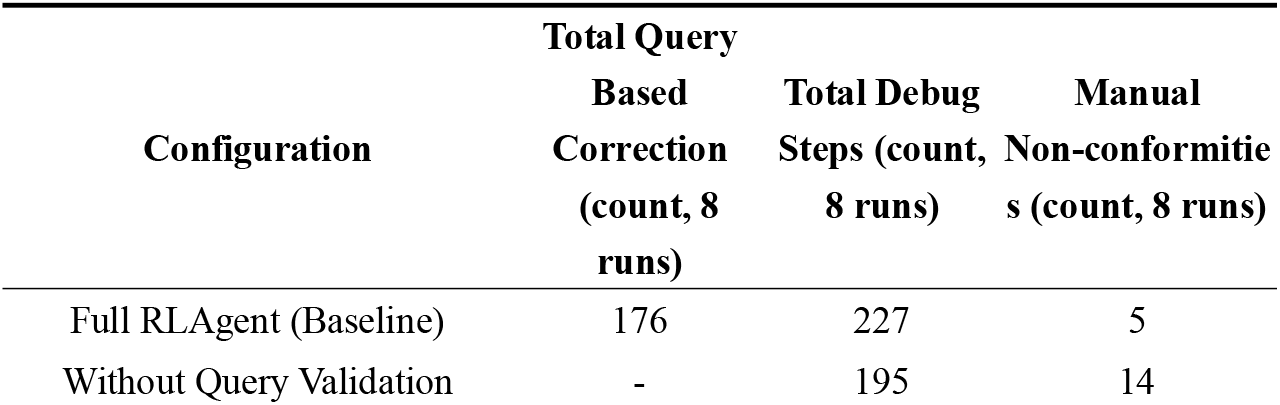
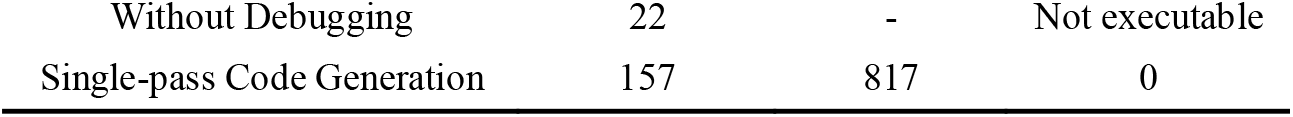
Ablation study of RLAgent components using DeepSeek-R1:70B as the backend. Each configuration was tested across eight full workflow runs.

The results highlight several findings. First, debugging steps were consistently more frequent than query-based correction, underscoring that error correction is a major component of the workflow’s computational cost. When query validation was removed, the total number of debugging iterations decreased, but at the expense of a higher number of manual non-conformities (14 across 8 runs), showing that query validation plays a central role in ensuring correctness. When debugging was removed, workflows could not be executed to completion, confirming that automated error correction is indispensable for system reliability.

Interestingly, the single-pass code generation strategy achieved perfect conformity, with zero manual corrections required. This suggests that providing the model with a complete specification in a single prompt may allow it to leverage contextual coherence more effectively, producing highly accurate code. However, this came at a substantial cost: query-based correction and debugging steps increased dramatically (157 and 817, respectively, across 8 runs), leading to significantly longer runtimes that could extend from hours to days.

Overall, these results highlight the complementary contributions of query validation and debugging to the robustness of RLAgent, while also revealing that stepwise generation offers a pragmatic balance between accuracy and efficiency. The contrast with single-pass generation suggests that future improvements may benefit from hybrid strategies, combining the contextual strength of complete specifications with the efficiency of modular, stepwise refinement.

### C. Architecture-Agnostic Performance of RLAgent Across Classical and Deep Models

RLAgent’s unified, language-driven pipeline adapts cleanly across heterogeneous model families (Table 3). Deep architectures achieved the strongest overall performance, with the Transformer yielding the top AUC (0.941) and F1-score (0.903), closely followed by Mamba (AUC 0.933, F1 0.886).

**Table 3.** Model performance under the RLAgent pipeline (with and without optional RAG)

| Model | ACC | Precision | Recall | F1-score | AUC |
| --- | --- | --- | --- | --- | --- |
| XGBoost | 0.8841 | 0.8866 | 0.8841 | 0.8846 | 0.9144 |
| XGBoost (RAG) | 0.6739 | 0.6790 | 0.6739 | 0.6756 | 0.7674 |
| Naïve Bayes | 0.5072 | 0.3979 | 0.5072 | 0.4229 | 0.4410 |
| Naïve Bayes (RAG) | 0.5000 | 0.4361 | 0.5000 | 0.4468 | 0.4607 |
| Random Forest | 0.8551 | 0.8600 | 0.8551 | 0.8559 | 0.9209 |
| Random Forest (RAG) | 0.6449 | 0.6652 | 0.6449 | 0.6475 | 0.7543 |
| lightGBM | 0.8623 | 0.8685 | 0.8623 | 0.8632 | 0.9179 |
| lightGBM (RAG) | 0.6377 | 0.6497 | 0.6377 | 0.6404 | 0.7332 |
| Transformer | 0.8913 | 0.9459 | 0.8642 | 0.9032 | 0.9409 |
| Transformer (RAG) | 0.6957 | 0.7010 | 0.8395 | 0.7640 | 0.7160 |
| LSTM | 0.8623 | 0.8690 | 0.9012 | 0.8848 | 0.8781 |
| LSTM (RAG) | 0.7319 | 0.7340 | 0.8519 | 0.7886 | 0.7559 |
| Mamba | 0.8696 | 0.9091 | 0.8642 | 0.8861 | 0.9333 |
| Mamba<br>(RAG) | 0.7536 | 0.7640 | 0.8395 | 0.8000 | 0.8399 |

Among classical learners, tree-based ensembles remained competitive (XGBoost AUC 0.914; Random Forest AUC 0.921; LightGBM AUC 0.918), underscoring that RLAgent’s standardized preprocessing and feature assembly produce compact descriptors that non-neural methods can exploit effectively. Notably, LSTM exhibited the highest recall (0.901) while trailing the Transformer in AUC, illustrating that RLAgent enables architecture selection aligned with application priorities (for example, high-recall screening versus higher-precision design).

As a secondary observation, adding RAG features in this small-sample setting generally reduced accuracy and AUC across models, and often shifted the precision-recall balance (for example, Transformer + RAG shows relatively higher sensitivity than precision). This suggests that external knowledge can act as noise when coverage is sparse or misaligned with region-level signals. Practically, RLAgent’s optional RAG should be activated conditionally (for larger, heterogeneous datasets or when coverage is verified), while the core takeaway here is architectural: RLAgent provides a model-agnostic harness that supports rapid, reproducible comparison and deployment across classical and deep learners.

### D. Case Study: Interactive Region-Level Prediction

To further illustrate RLAgent’s interactive modeling capabilities, we conducted a case study using the processed RNA-ligand dataset and selected the Mamba model for training (Fig. 3 and Fig. 4). After receiving task instructions from RLAgent, the user confirmed that the training data was prepared in the prescribed format. RLAgent automatically validated the input file, performed feature encoding, and initialized the training pipeline based on the user-specified model.

**Fig. 3.**
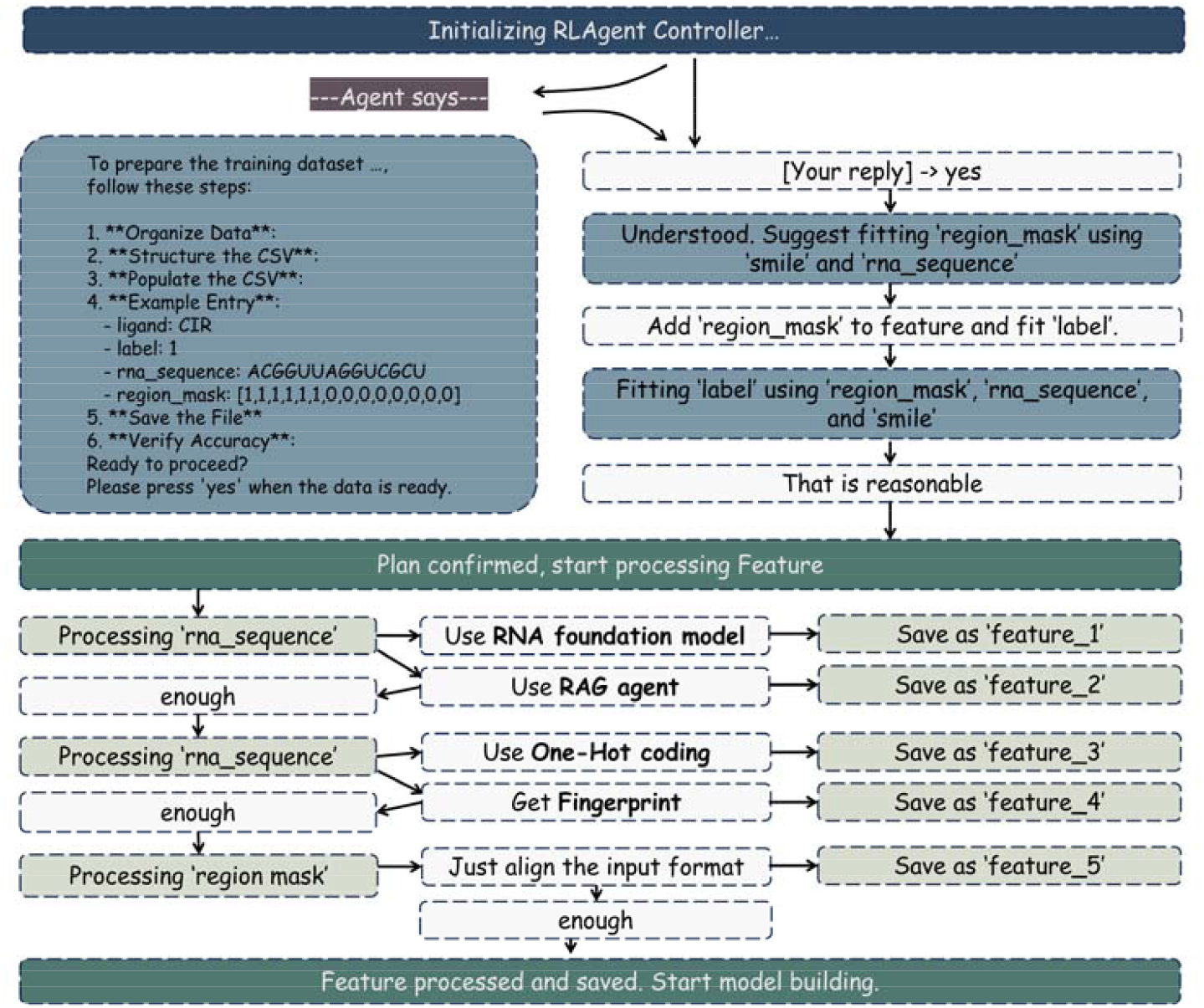
Dialogue-driven initialization and feature processing workflow of RLAgent: a case.

**Fig. 4.**
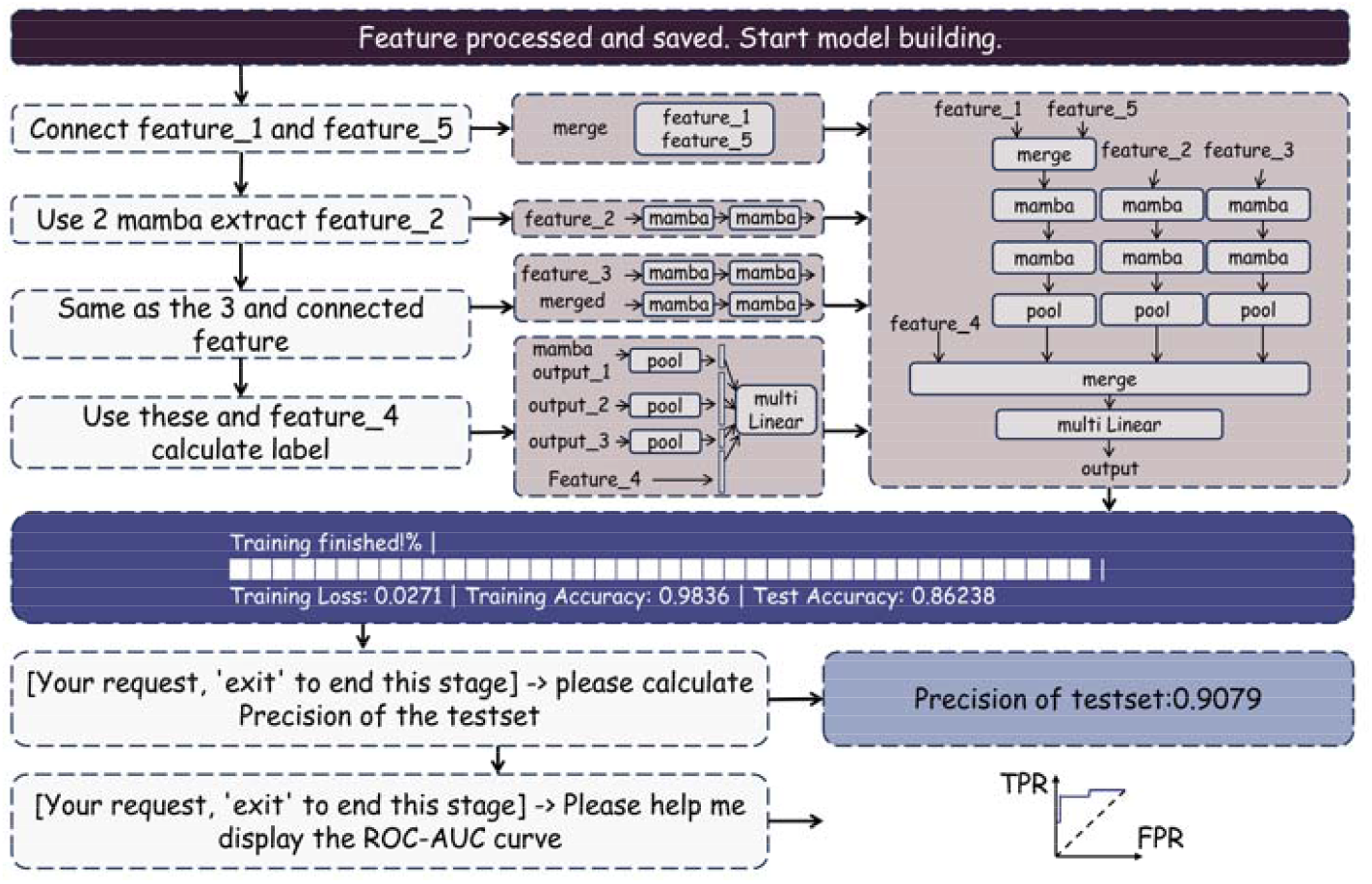
Model construction and evaluation workflow of RLAgent: a case.

During the workflow, all major configurations, including model selection, feature processing, and evaluation options, were specified through natural language commands rather than direct coding. RLAgent then executed the pipeline, reporting training progress, intermediate metrics, and debugging information when needed. The final outputs included quantitative results, such as accuracy on both training and test sets, and visual summaries, including the confusion matrix and ROC curve.

This case study demonstrates how RLAgent provides a seamless and user-friendly experience in region-level RNA-ligand prediction. By integrating natural language interaction with automated pipeline execution, RLAgent lowers technical barriers, ensures reproducibility, and delivers clear visualizations for model interpretation. These capabilities underscore the feasibility of applying RLAgent in real-world biomedical research, where accessibility, reliability, and interpretability are essential for effective computational modeling.

## IV Discussion

In this study, we introduced RLAgent, an interactive agent framework for modeling RNA region-ligand interactions. Unlike conventional tools that focus on global sequence or binding site predictions, RLAgent operates at the region level, offering a more flexible modeling perspective that accommodates complex binding patterns. By integrating local LLMs, RLAgent enables natural language-based control over the entire modeling pipeline, from feature encoding to model selection and visualization.

Despite these contributions, the current version of RLAgent has several limitations. The training dataset remains relatively small, which may introduce overfitting risks and limit generalizability. In addition, the weak supervision strategy for constructing negative samples may introduce false negatives, although such assumptions are common in early-stage molecular modeling. We emphasize, however, that RLAgent is designed primarily as a framework for enabling agent-based, modular, and code-free RNA-ligand modeling workflows. As more experimentally validated data becomes available, the framework can support larger-scale, statistically robust studies with broader biological impact. In future versions, RLAgent may be extended to support multi-resolution prediction, combining both region-level interaction modeling and fine-grained binding site identification.

In the longer term, RLAgent could serve as a general-purpose interface between domain experts and large RNA-ligand modeling backends, enabling human-AI collaborative hypothesis generation and accelerating RNA-targeted therapeutic design.

## V Conclusion

This study presented RLAgent as a new paradigm for interactive and region-level RNA-ligand modeling. Through robustness analysis, ablation experiments, model benchmarking, and a case study, we showed that RLAgent adapts to diverse architectures while highlighting the importance of model scale, workflow components, and external knowledge in predictive performance. With its automated debugging and iterative refinement, RLAgent provides reliable execution and suggests a potential path toward virtual experiments that can be interactively adjusted and improved.

More broadly, RLAgent illustrates how large language model-driven agents can contribute to computational biology by lowering technical barriers, enhancing reproducibility, and supporting transparent decision-making. By combining natural language interaction with automated workflows, RLAgent offers a reference framework for future applications of intelligent agents in RNA-targeted therapeutics and related areas.

## Competing interests

The authors declare no competing interests.

## Notes

### Competing Interest Statement

The authors have declared no competing interest.

### Summary of Updates

Title updated. Methods rewritten. Algorithm format updated.

## References

1. Zhou, Y., Y. Jiang, and S.J. Chen, RNA-ligand molecular docking: advances and challenges. Wiley Interdiscip Rev Comput Mol Sci, 2022. 12(3).

2. Zhuo, C., et al., Advances and Mechanisms of RNA-Ligand Interaction Predictions. Life (Basel), 2025. 15(1).

3. Kovachka, S., et al., Small molecule approaches to targeting RNA. Nat Rev Chem, 2024. 8(2): p. 120–135.

4. Morishita, E.C. and S. Nakamura, Recent applications of artificial intelligence in RNA-targeted small molecule drug discovery. Expert Opin Drug Discov, 2024. 19(4): p. 415–431.

5. Deigan, K.E. and A.R. Ferré-D’Amaré, Riboswitches: discovery of drugs that target bacterial gene-regulatory RNAs. Acc Chem Res, 2011. 44(12): p. 1329–38.

6. Haga, C.L. and D.G. Phinney, Strategies for targeting RNA with small molecule drugs. Expert Opin Drug Discov, 2023. 18(2): p. 135–147.

7. Yu, A.M., Y.H. Choi, and M.J. Tu, RNA Drugs and RNA Targets for Small Molecules: Principles, Progress, and Challenges. Pharmacol Rev, 2020. 72(4): p. 862–898.

8. Li, J., et al., Artificial intelligence for RNA-ligand interaction prediction: advances and prospects. Drug Discov Today, 2025. 30(6): p. 104366.

9. Morishita, E.C., Discovery of RNA-targeted small molecules through the merging of experimental and computational technologies. Expert Opin Drug Discov, 2023. 18(2): p. 207–226.

10. Fulle, S. and H. Gohlke, Molecular recognition of RNA: challenges for modelling interactions and plasticity. J Mol Recognit, 2010. 23(2): p. 220–31.

11. Wang, K., et al., RLBind: a deep learning method to predict RNA-ligand binding sites. Brief Bioinform, 2023. 24(1).

12. Wei, J., et al., Protein-RNA interaction prediction with deep learning: structure matters. Brief Bioinform, 2022. 23(1).

13. Sun, L.Z., et al., RLDOCK: A New Method for Predicting RNA-Ligand Interactions. J Chem Theory Comput, 2020. 16(11): p. 7173–7183.

14. Jiang, Y. and S.J. Chen, RLDOCK method for predicting RNA-small molecule binding modes. Methods, 2022. 197: p. 97–105.

15. Nithin, C., et al., Comparative analysis of RNA 3D structure prediction methods: towards enhanced modeling of RNA-ligand interactions. Nucleic Acids Res, 2024. 52(13): p. 7465–7486.

16. Lyu, M.R., et al., Automatic Programming: Large Language Models and Beyond. ACM Trans. Softw. Eng. Methodol., 2025. 34(5): p. Article 140.

17. Huang, H., et al., ProtChat: An AI Multi-Agent for Automated Protein Analysis Leveraging GPT-4 and Protein Language Model. J Chem Inf Model, 2025. 65(1): p. 62–70.

18. Schauperl, M. and R.A. Denny, AI-Based Protein Structure Prediction in Drug Discovery: Impacts and Challenges. J Chem Inf Model, 2022. 62(13): p. 3142–3156.

19. Ghafarollahi, A. and M.J. Buehler, ProtAgents: protein discovery via large language model multi-agent collaborations combining physics and machine learning. Digit Discov, 2024. 3(7): p. 1389–1409.

20. Inoue, Y., et al., DruGagent: Multi-Agent Large Language Model-Based Reasoning for Drug-Target Interaction Prediction. ArXiv, 2025.

21. Shen, T., et al., Accurate RNA 3D structure prediction using a language model-based deep learning approach. Nat Methods, 2024. 21(12): p. 2287–2298.

22. Sun, S., J. Yang, and Z. Zhang, RNALigands: a database and web server for RNA-ligand interactions. Rna, 2022. 28(2): p. 115–122.

23. Mapleson, D., et al., Efficient and accurate detection of splice junctions from RNA-seq with Portcullis. Gigascience, 2018. 7(12).

24. Hähnke, V.D., S. Kim, and E.E. Bolton, PubChem chemical structure standardization. J Cheminform, 2018. 10(1): p. 36.

25. Cavalleri, E., et al., An ontology-based knowledge graph for representing interactions involving RNA molecules. Scientific Data, 2024. 11(1): p. 906.

26. Gu, A. and T. Dao, Mamba: Linear-time sequence modeling with selective state spaces. arXiv preprint arXiv:2312.00752, 2023.

